# Interspecies Functional Divergence: The Microbiome’s Role in FLOT Response e Across Gastric Cancer Subtypes

**DOI:** 10.1101/2025.09.13.676018

**Authors:** Dionison Pereira Sarquis, Valéria Cristiane Santos da Silva, Daniel de Souza Avelar, Jéssica Manoelli Costa da Silva, Ronald Matheus da Silva Mourão, Taíssa Maíra Thomaz Araújo, Diego Pereira, Polyana Silva de Araújo, Geraldo Ishak, Samir Mansour Casseb, Samia Demachki, Williams Fernandes Barra, Fabiano Cordeiro Moreira, Rommel Mario Rodriguez Burbano, Paulo Pimentel de Assumpção

## Abstract

**Background:** Gastric adenocarcinoma exhibits marked molecular and histological heterogeneity, which is reflected in distinct patterns of progression, therapeutic response, and prognosis. Although the FLOT regimen (5-fluorouracil, leucovorin, oxaliplatin, and docetaxel) represents the current standard for perioperative chemotherapy, its systemic effects on the tumor microenvironment, including the associated bacterial microbiome and host gene expression, remain poorly understood.

**Methods:** This study investigated the effects of FLOT on the functional and ecological structure of intestinal and diffuse subtype gastric tumors by assessing its simultaneous influence on the human transcriptome, the bacterial transcriptome, and inter-kingdom interactions. We analyzed 55 tumor samples (37 intestinal subtype; 18 diffuse subtype) and explored potential genetic-functional interactions between the bacterial microbiome and the human genome.

**Results:** The results reveal a highly specific functional pattern in the diffuse subtype, absent in the intestinal subtype, demonstrating a unique ecological-transcriptional plasticity mediated by the microbiota under chemotherapeutic pressure. The interaction network was dominated by high-magnitude positive correlations. Notably, the bacterial gene *leuS* showed a robust association with the human gene *HCN1* and processes such as *potassium ion transmembrane transport*, *membrane depolarization*, and *regulation of postsynaptic membrane potential*, indicating a coordinated activation of ion channels and neuroepithelial circuits. Bacterial species including *Bacteroides uniformis*, *Faecalibacterium prausnitzii*, *Butyrivibrio crossotus*, *Prevotella copri*, and *Simiaoa sunii* also converged functionally on *HCN1*. Additionally, bacterial genes *mfd, nifJ, secY, rplF,* and *tet(Q)* were associated with pathways related to cell adhesion, epithelial proliferation, membrane potential control, and synaptic transduction.

**Discussion:** Integrative analysis reveals that the FLOT regimen acts as a systemic remodeler of the gastric tumor microenvironment, exerting distinct effects according to the histological subtype. While the intestinal subtype responds more aligned with the cytotoxic goals of chemotherapy, the diffuse subtype exhibits a functional plasticity that favors the emergence of adaptive and possibly pro-tumoral phenotypes. We propose a mechanistic model where in chemotherapy selectively reshapes the microbial ecosystem, which in turn modulates host functional circuits, directly influencing tumor behavior. These findings open perspectives for combined therapeutic strategies that include targeted modulation of the microbiome as an adjuvant to chemotherapy.

## Introduction

Gastric adenocarcinoma (GA) is a highly heterogeneous malignant neoplasm and represents one of the leading causes of cancer-related mortality worldwide^1,2^. Historically classified into intestinal and diffuse subtypes according to the Laurén classification, GA exhibits substantial differences in terms of tissue architecture, progression pathways, and therapeutic response^3,4^. This heterogeneity poses significant challenges to the standardization of clinical strategies and the development of personalized approaches.

In recent years, the FLOT regimen (5-fluorouracil, leucovorin, oxaliplatin, and docetaxel) has been established as the standard perioperative chemotherapy treatment for patients with locally advanced gastric tumors. However, investigations assessing the effects of FLOT beyond classical cytotoxicity remain scarce, particularly concerning the functional remodeling of the tumor microenvironment, the modulation of the human transcriptome, and alterations in the associated microbial ecosystem^5^.

Emerging evidence indicates that the tumor microbiome directly influences response to chemotherapy, whether through drug metabolism, local immune regulation, or the activation of inter-kingdom molecular axes. In parallel, studies have shown that bacterial and human genes can establish integrated functional circuits, phenotypically shaping the tumor under therapeutic stress^6,7,8^. Despite these advances, there is, to date, no integrated characterization of the transcriptomic, taxonomic, and functional responses induced by FLOT in the different GA subtypes.

In this context, this study proposes a multidimensional approach, evaluating how FLOT simultaneously reorganizes the human transcriptome, the tumor microbiome, and inter-kingdom molecular interactions in intestinal and diffuse subtype tumors. The central hypothesis is that the effects of FLOT are ecologically mediated and modulated by the pre-existing functional topology of each subtype, and are therefore divergent in impact, depth, and biological outcome. By integrating multi-scale data and robust statistical analyses, we seek to elucidate the chemotherapeutic action in gastric cancer, paving the way for a precision oncology framework based on tumor-microbiome-drug networks^9,10,11^.

## Materials and methods

### Ethics statement

This study was approved by the Ethics and Research Committee of João de Barros Barreto University Hospital (approval number: 47580121.9.0000.5634) and was conducted in accordance with the principles outlined in the Declaration of Helsinki. Participant recruitment and sample collection were carried out between July 2, 2022, and July 6, 2023. Before enrollment, all participants received detailed information about the study’s objectives, potential benefits, risks, and possible harms, ensuring a thorough understanding of the research. Written informed consent was voluntarily obtained from all participants prior to their inclusion in the study.

### Cohort and Sample Collection

A total of 55 fresh GA tumor samples were included, collected at the time of gastrectomy from patients treated at the Hospital Universitário João de Barros Barreto in Pará, Brazil. Patients were divided into two main groups: those treated with neoadjuvant chemotherapy (FLOT; n = 23) and those who underwent direct surgery (upfront surgery; n = 32). The samples were histologically classified into diffuse (n = 21) and intestinal (n = 34) subtypes according to the Lauren criteria.

### RNA Extraction and Sequencing

Approximately 30 mg of tumor tissue from each sample was macerated for RNA extraction using the TRIzol Reagent method (Thermo Fisher Scientific), following the manufacturer’s instructions. The total RNA was then assessed for integrity and concentration (ng/µL) using the Qubit 4 Fluorometer (Thermo Fisher Scientific). The established criterion for the RNA Integrity Number (RIN) was a value of ≥ 5. After verifying integrity (RIN ≥ 5), libraries were prepared using the *TruSeq Stranded Total RNA kit* (*Illumina®*) and sequenced on the *Illumina NextSeq 500®* platform with a *paired*-*end* strategy (2×75 bp).

### Microbial Taxonomic Profiling and Quantification of Bacterial and Human Genes

Microbiome species characterization was performed by applying *Kraken2* (v2.1.4)^12^ with the 16 GB trimmed *PlusPF* database for taxonomic classification. Bacterial gene quantification was based on pseudoalignment using *Salmon*^13^; the genomes identified by *Kraken2* were downloaded from *NCBI*^14^ and subsequently transformed into a database for transcriptome alignment. Human genes were quantified with *Salmon* and annotated using the *GENCODE* v43 reference database^15^.

### Differential Expression Analysis

Differential expression of human and bacterial genes between groups was assessed using the *DESeq2* package (v1.34.0)^16^, with a specific design for comparison between treatment and subtype. Genes with a |*log₂FC*| > 1 and an *FDR* (Benjamini-Hochberg) < 0.05 were considered significant. The analysis was conducted separately for the following contrasts: (i) intestinal with FLOT vs. without FLOT, (ii) diffuse with FLOT vs. without FLOT, (iii) diffuse with FLOT vs. intestinal with FLOT.

### Multivariate Segmentation (DAPC)

To investigate the separation of molecular profiles between groups, Discriminant Analysis of Principal Components (*DAPC*) was applied using the *adegenet* package (v2.1.10)^17^. The analysis was conducted based on the expression of human and bacterial genes, as well as on species composition. The analyzed groups were treated vs. untreated and the diffuse vs. intestinal subtypes.

### Functional Enrichment

Differentially expressed human genes were subjected to functional enrichment analysis using the *ClusterProfiler* package (v4.2.2)^18^, considering biological ontologies from *Gene Ontology (GO:BP)*^19^. The *org.Hs.eg.db* database^20^ was used, with a significance threshold of *p.adjust* < 0.05 (BH method).

### Overlap Tests (Hypergeometric)

The similarity between gene or species sets across groups was assessed using hypergeometric tests with *FDR* correction, employing the *phyper* function in R. Specific comparisons included the overlap of expressed genes between treated and untreated groups within the same subtype, and between FLOT-treated subtypes.

### Inter-Kingdom Correlation

Correlations between bacterial genes, human genes, and biological processes were assessed using *Spearman’s* correlation test, considering only correlations with |ρ| > 0.6. The analyses were visualized via correlograms (*corrplot* package^21^) and *Sankey* diagrams (*ggalluvial* package^22^), enabling the identification of microbiome–transcriptome–function interactive axes.

## Results

### Modulation of the Tumor Microbiome

We identified that the genes and species comprising the gastric tumor microbiome undergo relevant modulations both as a function of FLOT treatment and between the diffuse and intestinal histological subtypes. Furthermore, these alterations also manifest distinctly within each subtype, as evidenced by the comparison between treated and untreated diffuse samples.

### Molecular Discrimination

The segmentation analysis by *DAPC* revealed a clear separation between treated and untreated tumors based on bacterial gene and species profiles, as well as between the intestinal and diffuse subtypes. A similar pattern of discrimination was observed in human gene expression, demonstrating the method’s capacity to distinguish the groups based on transcriptomic and metatranscriptomic signatures. We observed a clear separation between groups based on bacterial gene profiles, microbial species, and human genes (Fig. 1, 2, and 3).

**Figure 1.**
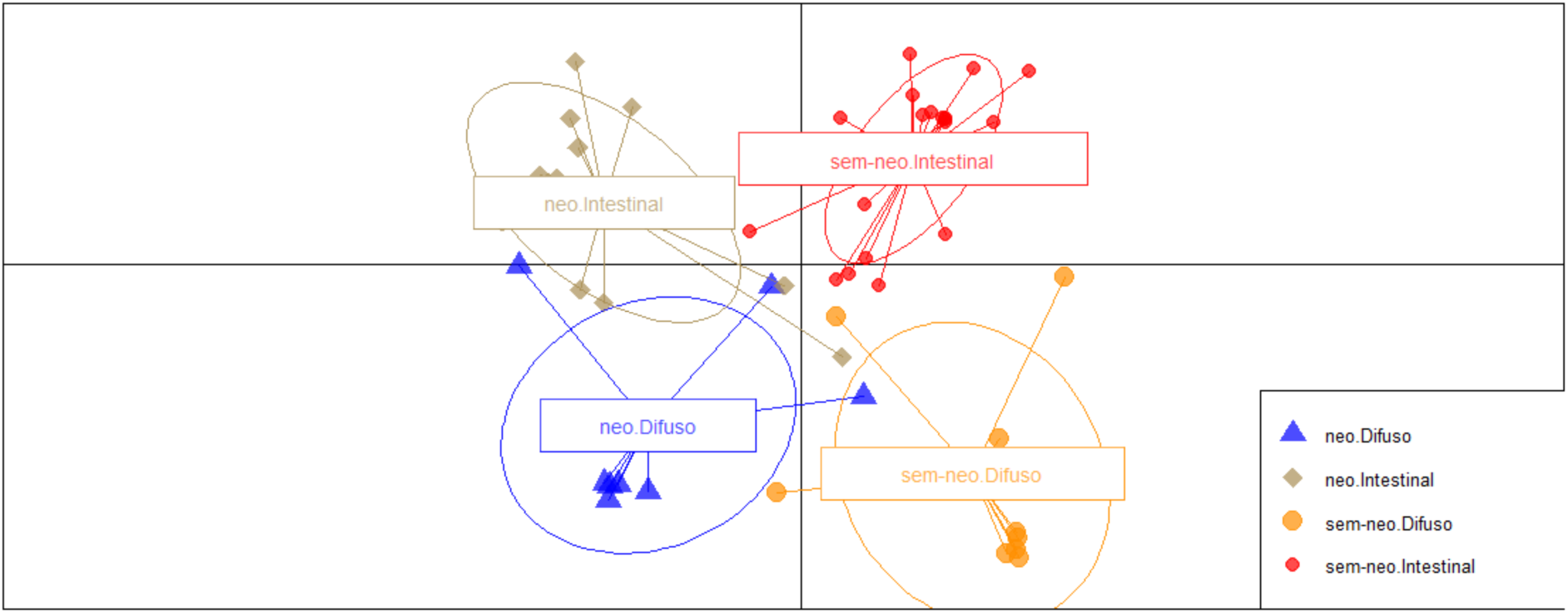
DAPC of microbiome genes across the samples.

**Figure 2.**
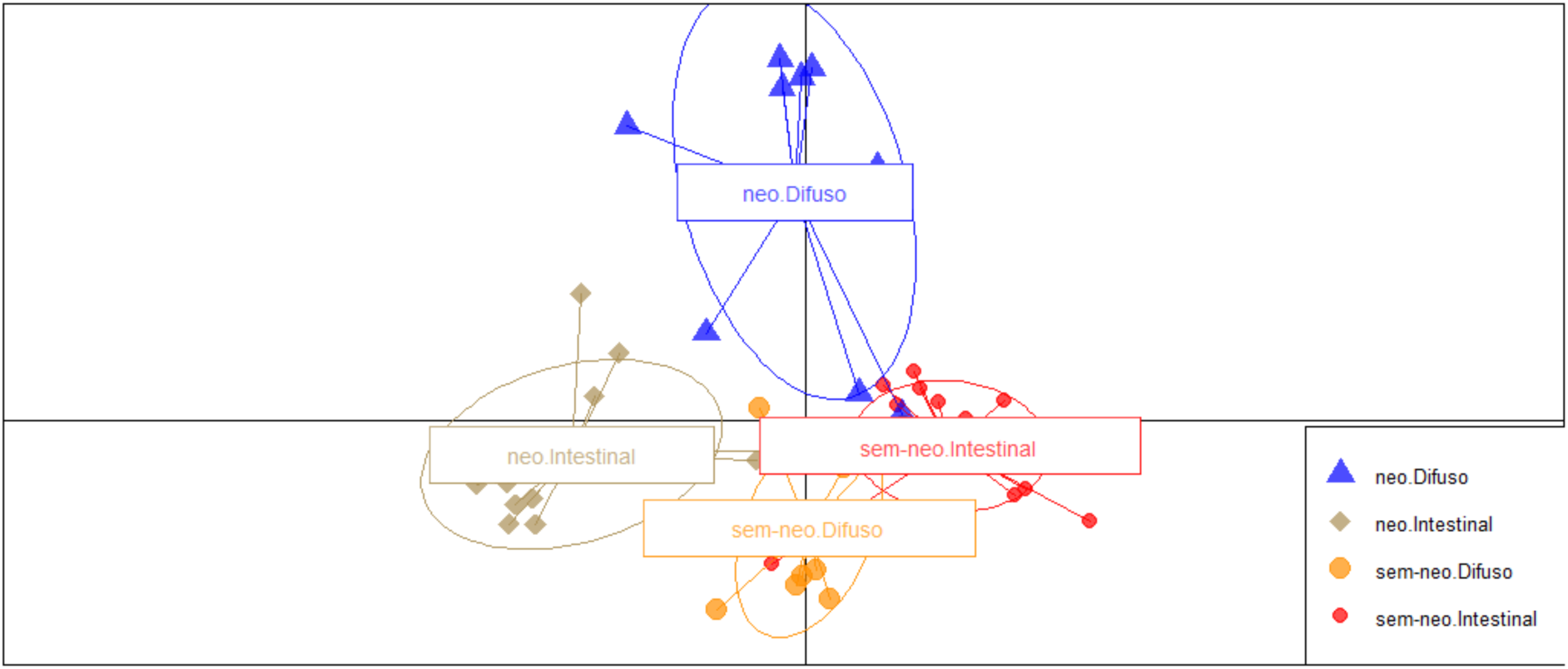
DAPC of microbiome species across the samples.

**Figure 3.**
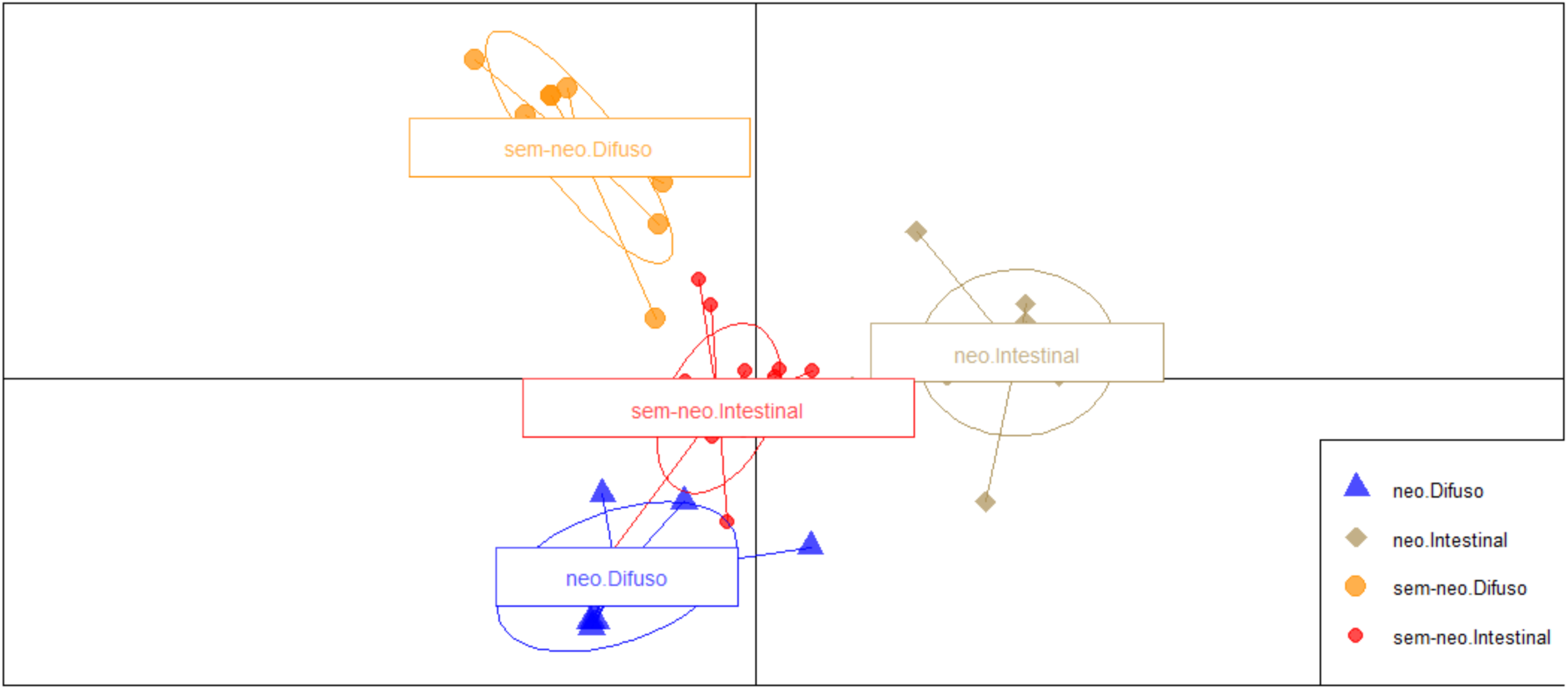
DAPC of human genes across the samples.

Furthermore, Venn diagram analysis revealed low overlap of genes and species between the compared groups (Fig. 4 and 5), reinforcing the specificity of the molecular signatures associated with each condition.

**Figure 4.**
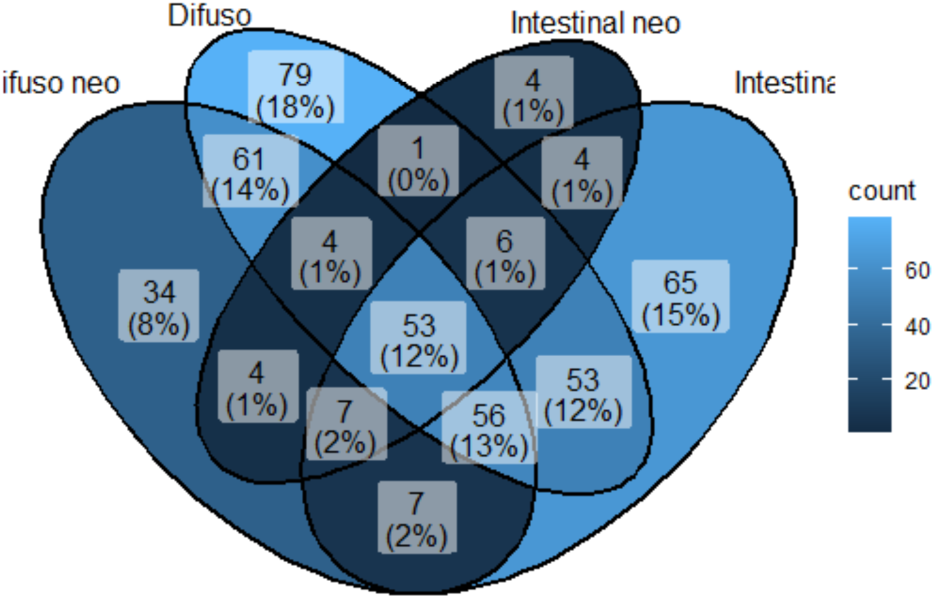
Venn diagram illustrating the overlap of microbiome genes across the samples.

**Figure 5.**
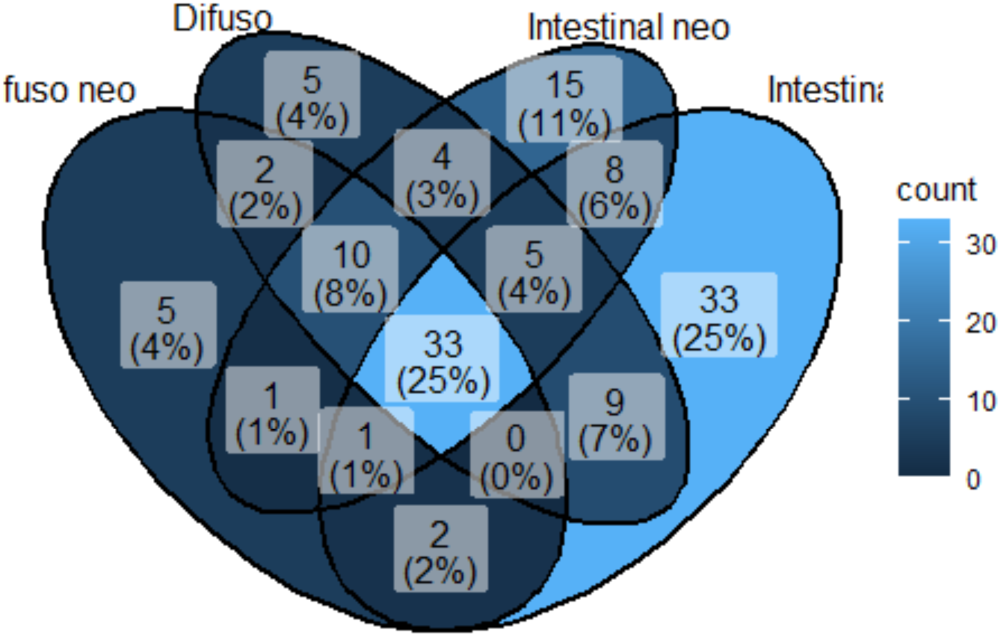
Venn diagram illustrating the shared and unique microbiome species across the samples.

Additionally, hypergeometric tests indicated statistically significant distinctions both as a function of treatment and between tumor subtypes (Tables 1 and 2).

**Table 1.** – Hypergeometric test results for microbiome gene data.

| Gene | Difuso Flot | Difuso | Intestinal Flot | Intestinal |
| --- | --- | --- | --- | --- |
| Difuso Flot | - | 3.665705e-141 | 9.770862e-85 | 1.907068e-94 |
| Difuso | 3.665705e-141 | - | 6.645801e-72 | 1.615153e-130 |
| Intestinal Flot | 9.770862e-85 | 6.645801e-72 | - | 1.149132e-84 |
| Intestinal | 1.907068e-94 | 1.615153e-130 | 1.149132e-84 | - |

**Table 2.** – Hypergeometric test results for microbiome species data.

| Specie | Difuso Flot | Difuso | Intestinal Flot | Intestinal |
| --- | --- | --- | --- | --- |
| Difuso Flot | - | 2.2899e-72 | 3.569445e-67 | 1.018323e-43 |
| Difuso | 2.2899e-72 | - | 2.910791e-75 | 8.810924e-30 |
| Intestinal Flot | 3.569445e-67 | 2.910791e-75 | - | 4.285898e-27 |
| Intestinal | 1.018323e-43 | 8.810924e-30 | 4.285898e-27 | - |

We identified the main biological processes associated with each subtype and treatment condition. In the intestinal subtype without FLOT, processes related to extracellular matrix organization were prominent, including *extracellular matrix organization, extracellular structure organization*, and *external encapsulating structure organization* (Fig. 6). In the intestinal subtype with FLOT, functional processes focused on sensory perception and ionic signaling were observed, such as *detection of chemical stimulus involved in sensory perception, sensory perception of smell,* and *regulation of membrane potential* (Fig. 6). In the diffuse subtype with FLOT, the main processes involved epithelial proliferation and calcium-mediated signaling, including *epithelial cell proliferation, regulation of epithelial cell proliferation,* and *calcium-mediated signaling* (Fig. 7). It is important to note that no biological processes with statistical significance were identified in the diffuse subtype without FLOT.

**Figure 6.**
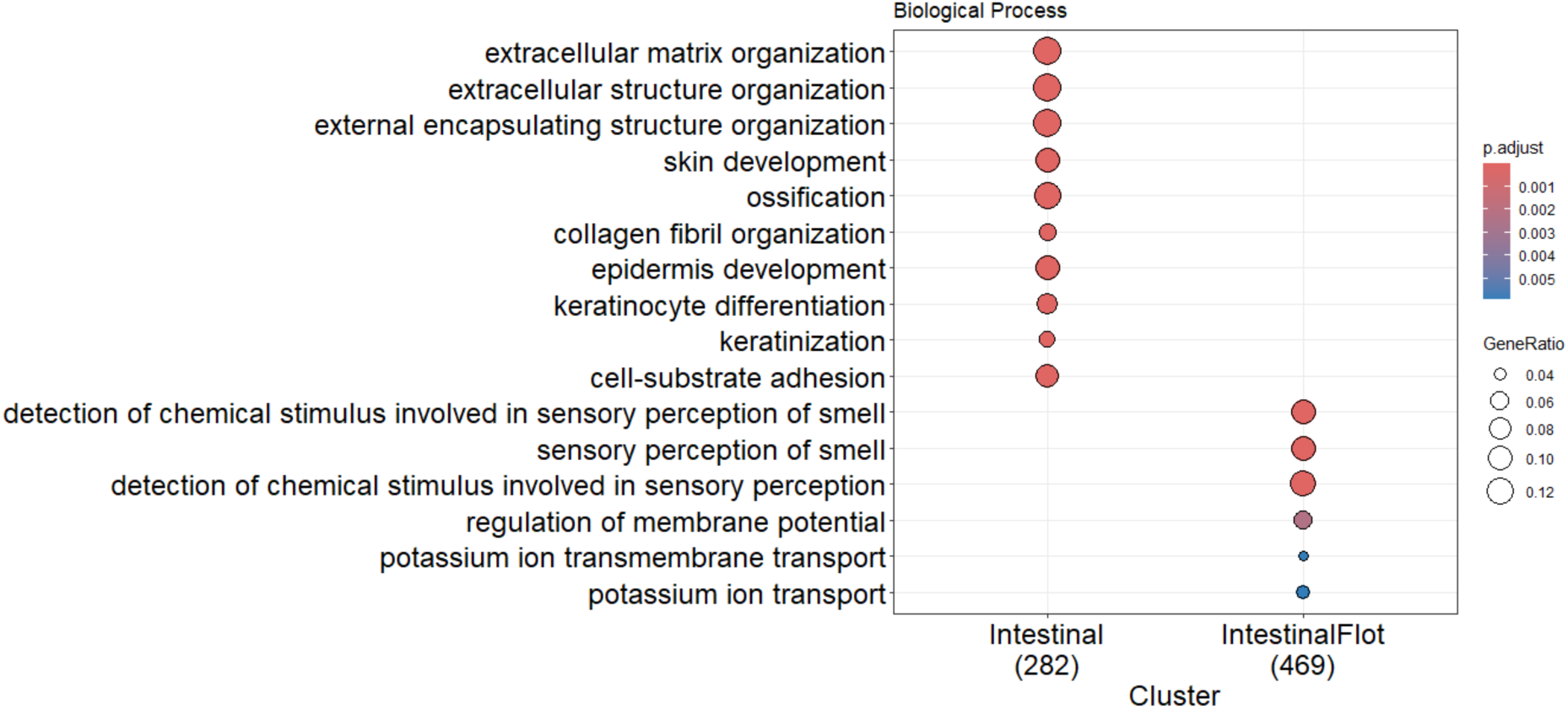
Comparison of intestinal biological processes with and without FLOT treatment.

**Figure 7.**
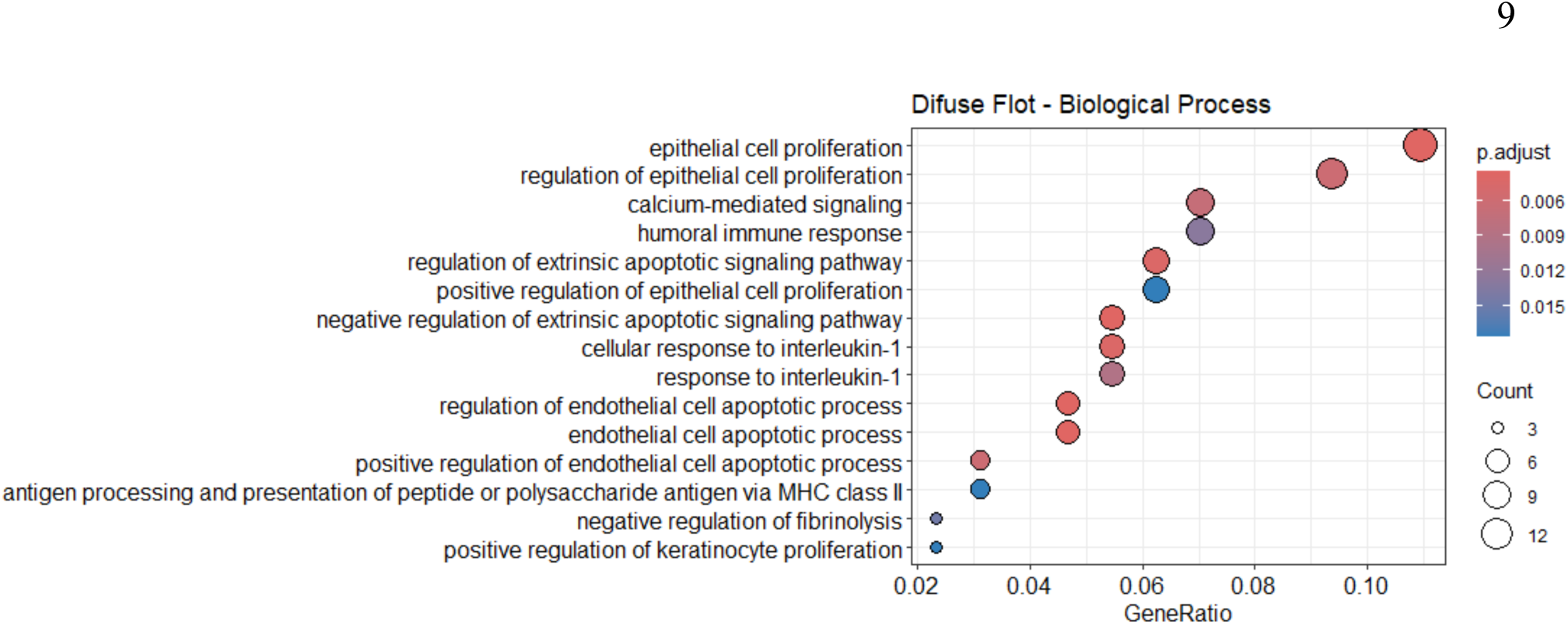
Biological processes in diffuse subtype with FLOT treatment.

### Human and Bacterial Gene Expression

We identified variation in human gene expression associated with treatment and histological subtype, mirroring the pattern observed in bacterial expression. Furthermore, both undergo distinct modulations depending on the interaction between the therapeutic regimen and the histological subtype. Additionally, we identified correlations between human transcriptomic and bacterial metatranscriptomic profiles, which are associated with biologically relevant processes in the tumor context.

### Diffuse Subtype Specifics

Regarding the microbiome, we observed a significant increase in the expression of bacterial genes and in the abundance of microbial species in FLOT-treated samples compared to untreated ones. Among the most highly expressed bacterial genes were *ilvD*, *pepN*, *sucB, mshA*, and *rpoD*. The enriched species included *Enterobacter cloacae*, *Enterobacter cloacae complex sp*., *Enterobacter kobei*, *Klebsiella oxytoca*, *Klebsiella variicola*, *Salmonella bongori, Streptococcus salivarius, Moraxella osloensis, Staphylococcus haemolyticus,* and *Streptococcus sp*.

Regarding the human component, the most relevant differentially expressed genes included *AREG, CCL4, CCR7, CHRNA7, ENKUR, GBP1, HAMP, HGF, IFNAR1, IGF1, IL1R2/IL1RN, MCL1, MMP2, OSM, REG1B, SELP, SERPINE1, THBS1,* and *WFDC3.* Functional analysis revealed significant enrichment in the following biological processes: *positive regulation of cell activation, regulation of epithelial cell proliferation, regulation of phosphatidylinositol 3-kinase/protein kinase B signal transduction, response to interleukin-1, humoral immune response, regulation of apoptotic signaling pathway, antigen processing* and *presentation, and receptor signaling pathway via JAK-STAT.* The correlations between microbial genes, human genes, and their respective biological processes are detailed in figures 8 and 9.

**Figure 8.**
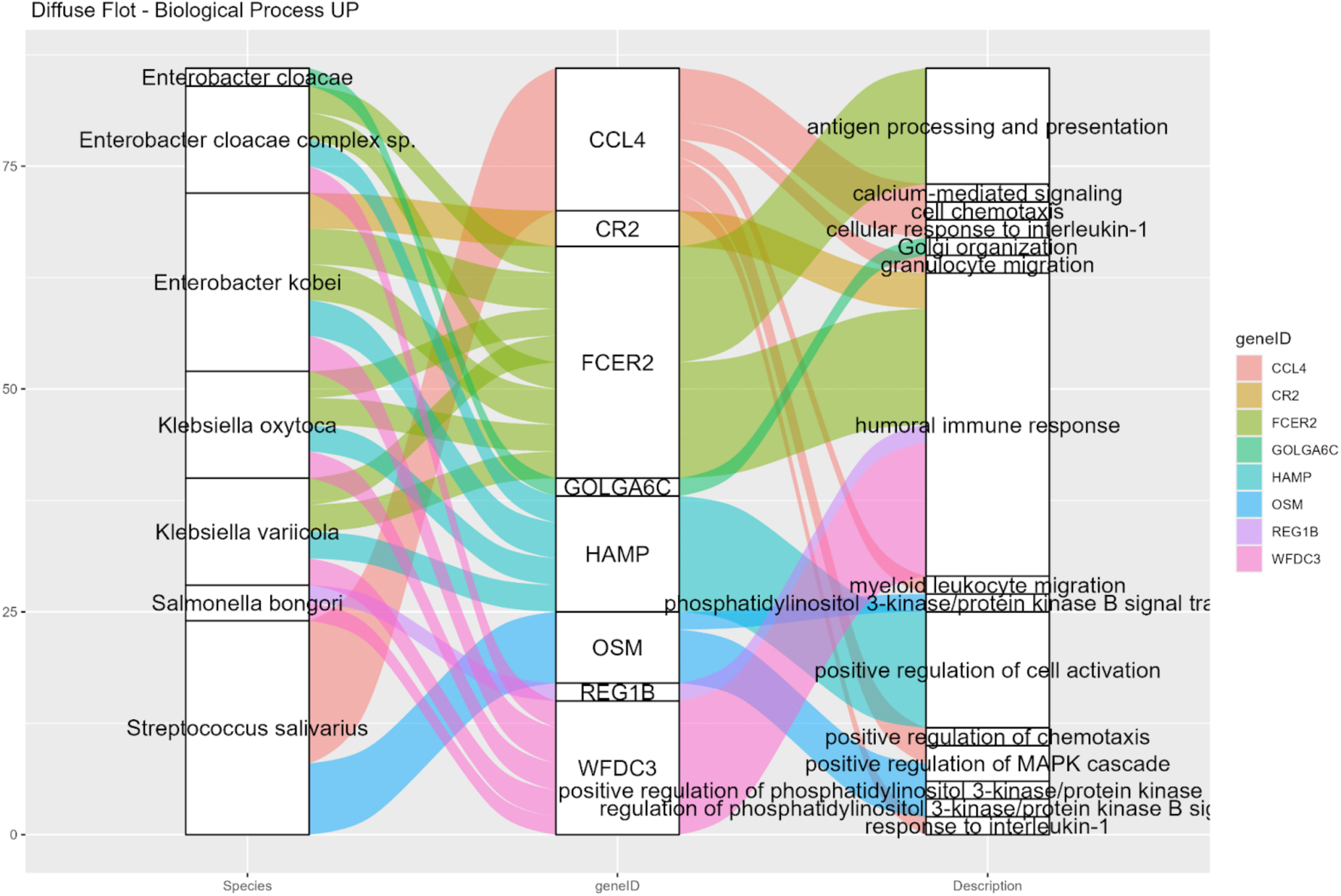
Sankey diagram of biological processes in the diffuse subtype with FLOT treatment, presenting correlations between human transcriptomic and bacterial metatranscriptomic profiles related to tumor-associated biological processes.

**Figure 9.**
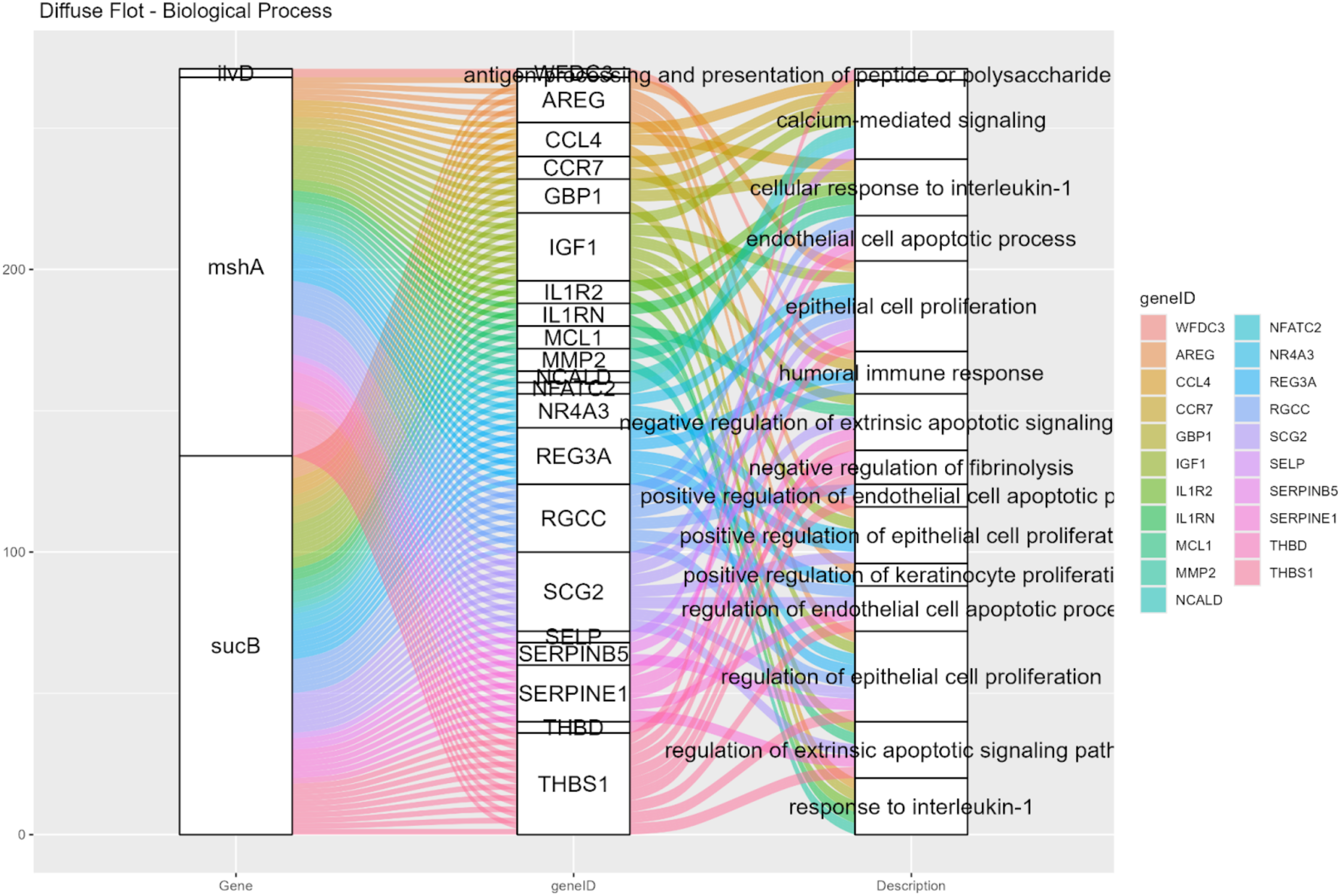
Sankey diagram of biological processes in the diffuse subtype with FLOT treatment, presenting correlations between human transcriptomic and bacterial gene expression related to tumor-associated biological processes.

### Intestinal Subtype Specifics

In the intestinal subtype not subjected to treatment, we observed enrichment of the bacterial genes *atpD, grpE, and ureB*, along with the species *Gemella haemolysans, Lactobacillus crispatus, Lactobacillus johnsonii, Porphyromonas sp. oral taxon 275, Prevotella intermedia,* and *Gemmata massiliana*. In the human component, the highly expressed genes *CXCL8, DLX6, FOSL1, GFAP, HOXC10, IL1B, KRT13, METRN, MUC21, CLDN6, CYP2W1,* and *GJB6* were prominent. The main enriched biological processes included *positive regulation of nervous system development, Regulation of epithelial cell proliferation, Inflammatory response / response to cytokine stimulus, epithelial differentiation,* and *response to lipopolysaccharide*.

In the treated intestinal subtype, we observed enrichment of the bacterial genes *zipA, hydB,* and *surA*, along with the species *Bacillus velezensis, Bacteroides uniformis, Butyrivibrio crossotus, Escherichia albertii, Klebsiella quasipneumoniae, Leptotrichia trevisanii, Micromonospora siamensis, Nocardia wallacei, Prevotella copri, Rhodococcus sp. 75, Simiaoa sunii, Staphylococcus haemolyticus, Streptomyces lydicus, Streptomyces sp. YPW6,* and *Bacillus cereus*. In the human transcriptome, genes such as *ABCC8, GRIN2B, HCN1, KCNB1, KCNA7, KCNJ11, KCNMB1, KCNMB4, NOS1, OR10A3, OR10H5, PRKN, RGS7,* and *SCN11A* were highlighted as highly expressed. The biological processes enriched in this group are *sensory perception of smell, regulation of membrane potential, potassium ion transport, potassium ion transmembrane transport, detection of chemical stimuli involved in sensory perception of smell,* and *detection of chemical stimuli involved in sensory perception.* The correlations between microbial genes, human genes, and their respective biological processes are detailed in figures 10 to 12.

**Figure 10.**
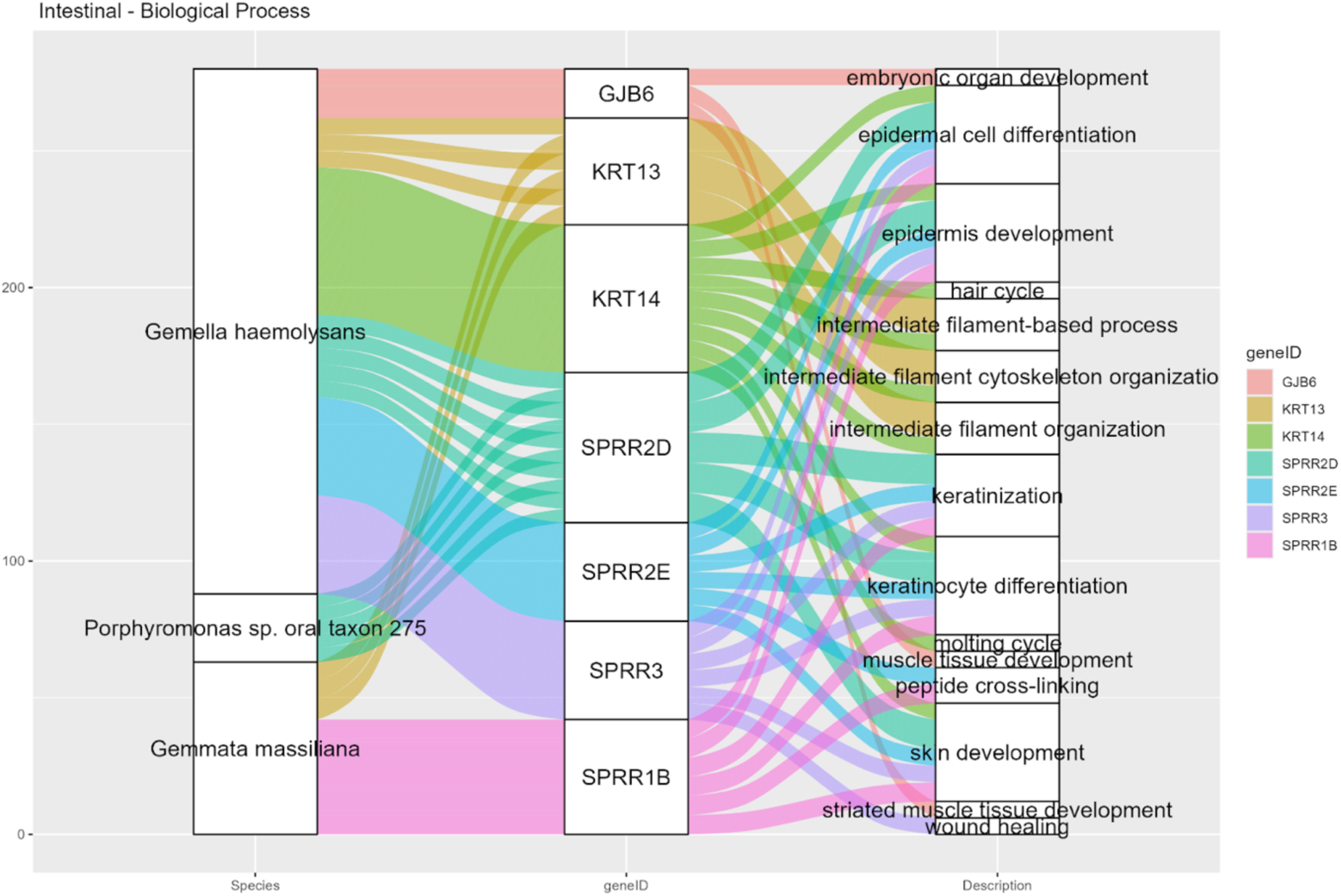
Sankey diagram of biological processes in the intestinal subtype without FLOT treatment, presenting correlations between human transcriptomic and bacterial metatranscriptomic profile related to tumor-associated biological processes.

**Figure 11.**
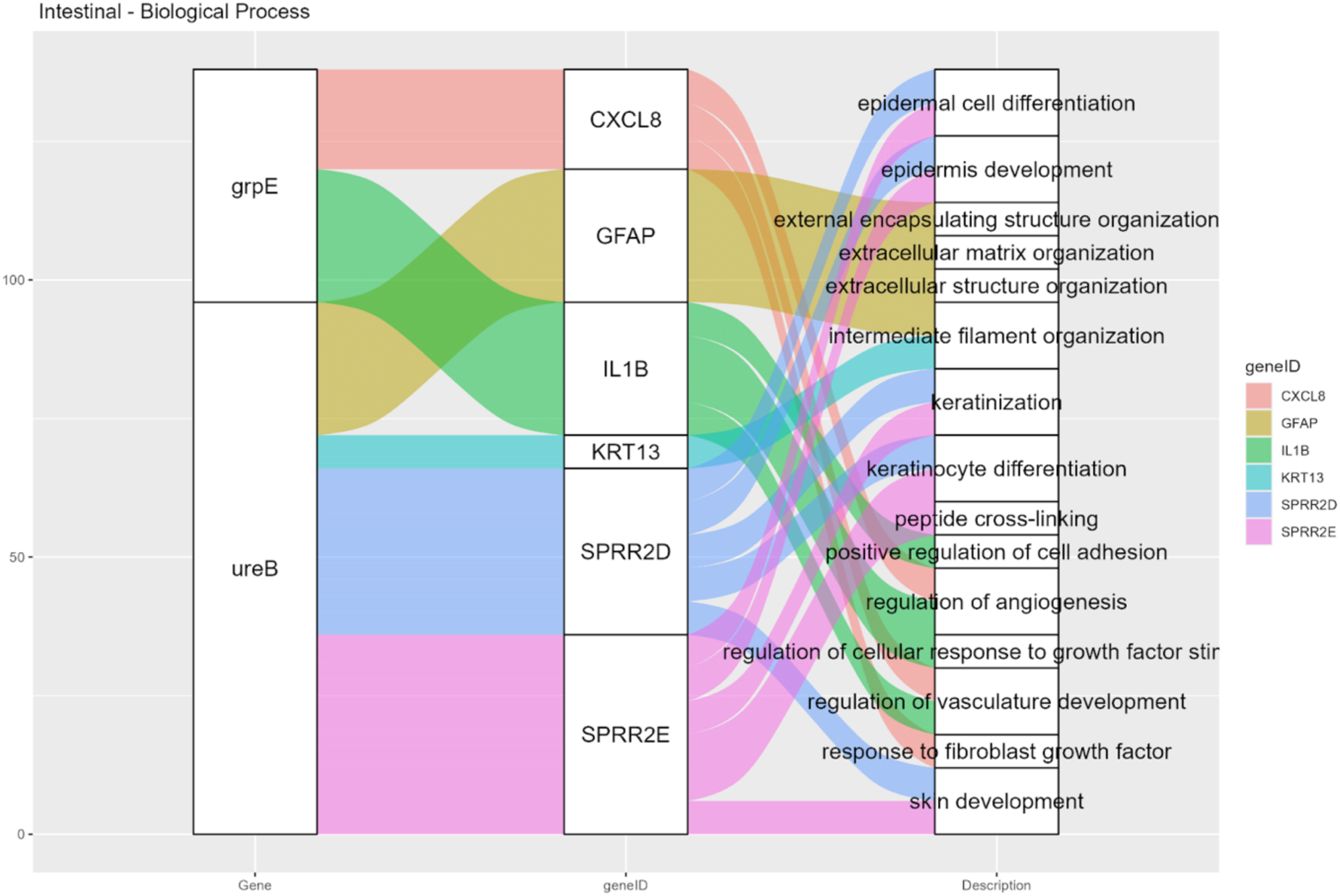
Sankey diagram of biological processes in the intestinal subtype without FLOT treatment, presenting correlations between human transcriptomic and bacterial gene expression related to tumor-associated biological processes.

**Figure 12.**
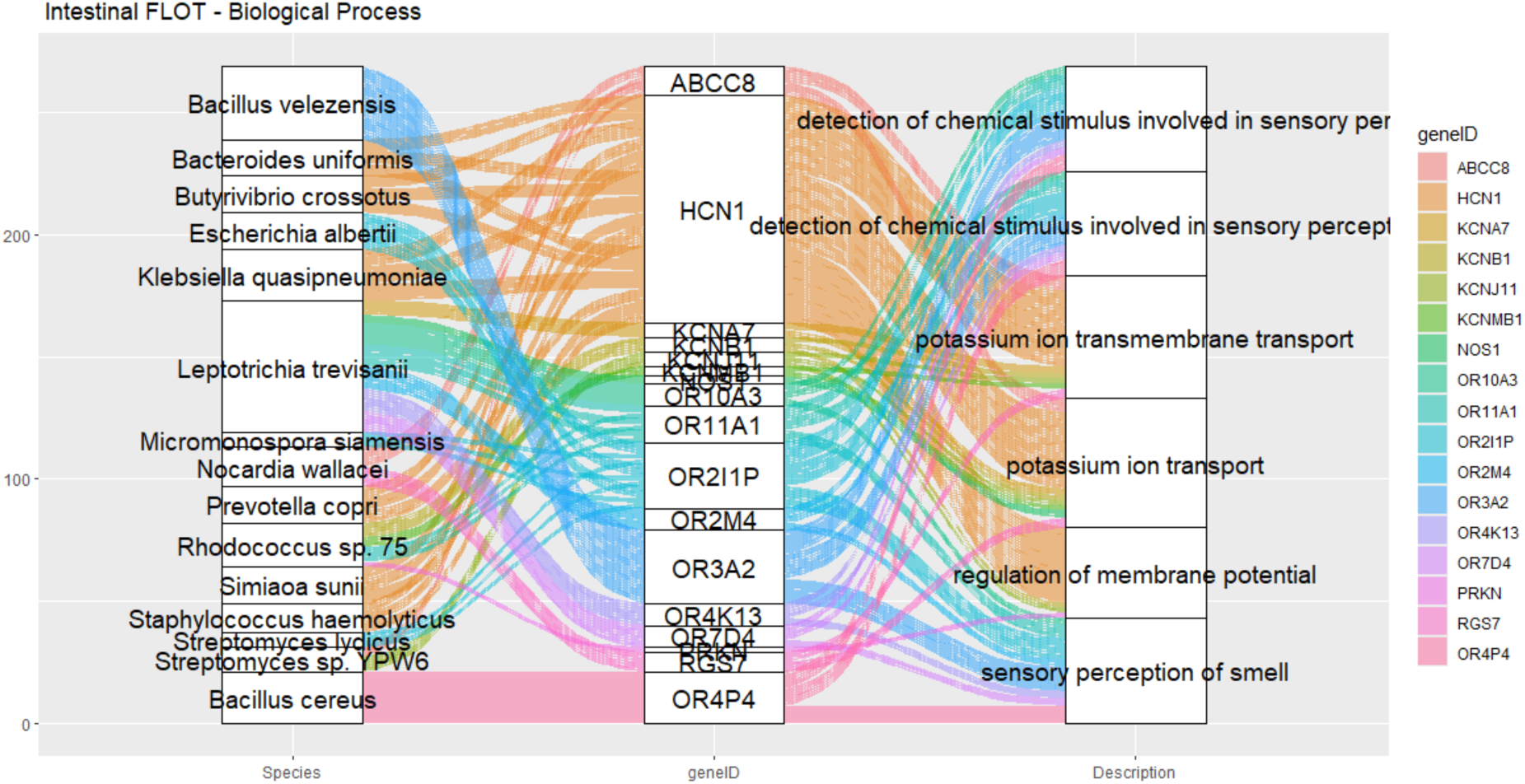
Sankey diagram of biological processes in the intestinal subtype with FLOT treatment, presenting correlations between human transcriptomic and bacterial metatranscriptomic profile related to tumor-associated biological processes.

## Discussion

This study reveals that the neoadjuvant FLOT regimen does not act solely as a cytotoxic protocol but rather as a complex modeling agent, with action on three fundamental levels of the gastric tumor microenvironment: (i) the human transcriptome; (ii) the composition and functionality of the microbiome; and (iii) the inter-kingdom network modulating the involved biological processes. This remodeling is highly dependent on the histological subtype of the tumor and reflects different degrees of biological plasticity, inflammation, and regenerative capacity.

Analysis of the intestinal subtype of gastric adenocarcinoma (GA) not subjected to FLOT treatment revealed a complex network of interactions between bacterial species, microbial genes, and human genes. Among the microbial genes, *atpD* exhibited a consistent association with the human genes *DLX6, FOSL1, HOXC10,* and *METRN.* These human genes are strongly implicated in processes related to cell differentiation, inflammation, and tissue development^23^. *FOSL1*, for example, is a transcription factor associated with inflammatory response and epithelial remodeling, while *DLX6* is involved in epithelial morphogenesis and the development of specific cell lineages^23^. The association of *atpD* with these genes suggests a functional role in activating pathways that favor tumor plasticity and structural reorganization of the microenvironment.

Complementarily, the bacterial gene grpE was notable for modulating the human genes *CXCL8, FOSL1, IL1B,* and *METRN*. The simultaneous presence of *CXCL8* and *IL1B* among the correlated genes indicates a coordinated activation of pro-inflammatory pathways in the tumor microenvironment, possibly sustained by continuous bacterial stimuli^24^. The correlation between *grpE* and *FOSL1*, shared with *atpD*, reinforces the hypothesis that bacterial products are directly associated with the transcription of human genes modulating immune response and epithelial proliferation^25^. Furthermore, the presence of *METRN*, a cytokine involved in angiogenesis and neuroprotection, as a common node between *atpD* and *grpE*, suggests a potential interface between neural and inflammatory processes mediated by bacterial stimuli^26^.

For ureB, positive correlations were observed with *GFAP, KRT13, MUC21, SPRR2D,* and *SPRR2E*. These genes are predominantly associated with epithelial differentiation and the maintenance of mucosal barrier integrity^27,28,29^. The coordinated expression of keratins and SPRR family proteins suggests a highly differentiated epithelial phenotype, associated with the formation of specialized barriers and resistance to environmental stresses^28^. In turn, MUC21, a transmembrane mucin, is implicated in the regulation of cellular adhesiveness and the protection of the epithelial surface, potentially also modulating interactions with the immune system and tumor progression^28,30^. Thus, the correlation pattern associated with ureB suggests that this bacterial gene may act as an indirect modulator of functionally altered epithelial differentiation, contributing to a microenvironment conducive to neoplastic progression.

The enriched biological processes, *keratinocyte differentiation, intermediate filament cytoskeleton organization, epidermis development, response to lipopolysaccharide, positive regulation of nervous system development, regulation of epithelial cell proliferation,* and *inflammatory response*, associated with the human genes (*DLX6, FOSL1, HOXC10, METRN, CXCL8, IL1B, GFAP, KRT13, MUC21, SPRR2D,* and *SPRR2E*) were positively correlated with the bacterial genes (*atpD, grpE,* and *ureB*). These data reinforce the notion that the microbiome actively participates in the functional orchestration of tumor tissue, modulating both its structure and immune response.

In the intestinal subtype subjected to neoadjuvant FLOT treatment, a complex reconfiguration of interactions between the microbiome, the human transcriptome, and critical biological processes was observed. Data integration revealed a robust correlation pattern between bacterial genes, particularly *zipA*, *hydB*, and *surA*, and human genes associated with electrochemical signaling, sensory perception, and ionic homeostasis, delineating a functional communication axis within the tumor microenvironment.

The bacterial gene *surA*, a periplasmic chaperone essential for the folding and assembly of outer membrane proteins in Gram-negative bacteria, exhibited a significant negative correlation with the human genes *CA6, HTR3B,* and *NEDD4. CA6* (Carbonic Anhydrase VI), which is involved in extracellular *pH* regulation and ion metabolism in secretory tissues, when suppressed, suggests that *surA* activity compromises acid-base homeostasis, promoting acidification of the tumor microenvironment—a condition widely associated with neoplastic progression, immune evasion, and therapy resistance^31,32^

In turn, *HTR3B*, a subunit of the 5-HT3 serotonin receptor, participates in neuroepithelial pathways related to nociception, intestinal motility, and paracrine signaling. Its repression indicates a potential deactivation of sensory circuits, rendering the tumor less responsive to environmental stimuli^33^. Meanwhile, *NEDD4*, an E3 ubiquitin ligase, regulates protein stability, cell adhesion, and stress response, playing a fundamental role in the functional integrity of the epithelium^34.35^. Its inhibition reinforces the idea that *surA* acts as a negative modulator of tumor resilience against environmental and pharmacological insults.

The bacterial gene *hydB*, associated with energy generation via hydrogenases, demonstrated a significant negative correlation with the human genes *RGS7, CNR2,* and *SCN11A. RGS7* is involved in modulating *G* protein-coupled receptors (*GPCRs*), regulating the amplitude and duration of extracellular signals^36^; *CNR2*, the type 2 cannabinoid receptor, has a recognized immunomodulatory and anti-inflammatory role, often explored in therapeutic strategies for tumor control^37,38^; and *SCN11A*, a voltage-gated sodium channel, acts in the regulation of membrane potential and the perception of nociceptive stimuli—functions essential for neuroepithelial signaling^39^.

Another highlight is the gene *ABCC8*, which encodes an *ATP*-sensitive potassium channel and is a member of the *ABC* transporter family. Its significant correlation with *Bacillus cereus* suggests a potential reconfiguration of tumor ion metabolism induced by this species. Although better known for its role in regulating insulin secretion, *ABCC8* has been implicated in drug resistance and the control of cellular excitability in tumor contexts, including gastric cancer^40,41^.

The integrative analysis of data from the diffuse subtype of gastric cancer treated with the FLOT regimen reveals a profound and coordinated reconfiguration between the tumor transcriptome and the associated microbial ecosystem. Beyond isolated gene expression changes, the findings demonstrate a complex functional architecture marked by molecular interactions among bacterial genes, microbial species, and human genes strongly involved in biological processes critical for tumor progression and therapeutic response.

Among the most expressive bacterial genes, *ilvD, pepN,* and *rpoD* stand out, whose positive correlations with human genes such as *HAMP, ENKUR,* and *GOLGA6C* point to a functional axis of immune activation, epithelial organization, and stress response. Notably, the association between *ilvD* and *HAMP* suggests activation of the *JAK*-*STAT* pathway, as well as engagement of iron metabolism regulation and leukocyte activation processes, key factors for tumor containment and the orchestration of adaptive immunity. The *pepN* gene, by correlating with *HAMP*, indicates a potential bacterial influence on secretion processes and cellular signaling, favoring the functional reprogramming of the tumor epithelium. Similarly, *rpoD*, a bacterial sigma factor essential for transcription, shows a positive association with *ENKUR* and *HAMP*, reinforcing the microbiome’s role in activating epithelial responses to chemotherapeutic stress^42,43,44,45,46^.

In contrast, *mshA* demonstrated marked negative correlations with genes such as *AREG* and *CCL4*, both strongly implicated in cell proliferation, inflammation, and epithelial migration^47,48^. This suppression may signal an inhibitory role of the microbiome on excessive tumor plasticity, potentially favoring a more orderly and less aggressive reprogramming of the microenvironment.

At the level of bacterial species, a set of taxa strongly correlated with human genes and core biological processes was evidenced. Species such as *Enterobacter cloacae complex, Streptococcus salivarius, Klebsiella variicola, Salmonella bongori, Staphylococcus haemolyticus,* and *Moraxella osloensis* exhibited robust correlations, both positive and negative, with human genes regulating cell motility, immune response, and intracellular signaling. A relevant example is the positive association of the *E*. *cloacae complex* with the process *cilium-dependent cell motility*, suggesting a direct influence on the migratory capacity of epithelial cells and, by extension, their invasive potential^49,50^

Conversely, species such as *Klebsiella variicola* and *Moraxella osloensis* exhibited a negative correlation with the process “*response to interleukin-1*”, a key pro-inflammatory axis^51^. This inhibition may represent a tumor adaptation to escape immune surveillance during treatment. Similarly, *Klebsiella oxytoca* showed a negative association with *regulation of apoptotic signaling pathways*, suggesting direct interference with the activation of programmed cell death pathways, a fundamental mechanism of tumor containment^52^.

Another significant finding was the correlation between *Salmonella bongori* and *processes such as insulin secretion* and *regulation of cell cycle G2/M phase transition*, indicating that the microbiome may modulate both the metabolism and the proliferation dynamics of tumor cells^53,54^. This suggests an adaptive role of certain bacterial species in sustaining mitotic activity and tumor plasticity even under chemotherapeutic pressure.

At the functional level, three major biological axes modulated by microbial interactions stand out: Adaptive Immunity and Antigen Processing, evidenced by *processes such as tumoral immune response, antigen processing* and *presentation and positive regulation of leukocyte activation*, suggests that part of the post-FLOT microbiome contributes to the recruitment and activation of immune system components, potentially enhancing or modulating their action^55^; Epithelial Regeneration and Structure, with the induction of processes such as *epithelial cell proliferation* and *keratinocyte proliferation*, indicates a dual role of the microbiome in mucosal restoration and, possibly, in sustaining proliferative niches^56^; Cellular Survival and Stress Resistance, with the activation of the *receptor signaling pathway via JAK-STAT*, combined with the absence of pro-apoptotic signaling, reveals a molecular environment more permissive to tumor survival and less responsive to cell death mechanisms, favoring therapeutic resistance^57^.

The integrative analysis of correlations between bacterial genes, bacterial species, human genes, and biological processes in the FLOT-treated diffuse subtype revealed a highly reorganized functional landscape distinct from the intestinal subtype, in which no enriched biological processes were identified. This dichotomy demonstrates that the effects of neoadjuvant FLOT treatment are modulated in markedly different ways according to the molecular and microbial context of each histological subtype, with the diffuse subtype being more susceptible to inter-species reprogramming.

The emerging functional architecture in the FLOT-treated diffuse subtype is dominated by high-magnitude positive correlations (r > 0.6), suggesting that the microbiome not only resists chemotherapy but is co-opted or favored by it, becoming functionally active in modulating the tumor epithelium. In this scenario, biological processes related to neuroplasticity, synaptic signaling, hormonal metabolism, and electrophysiological control stand out^58,59,60,61^.

On the bacterial side, genes such as *leuS, nifJ, mfd, rplF, ssrA, secDF, secY,* and *tet(Q)* were strongly correlated with critical human genes including *HCN1, DPYSL5, PCSK5, FGL1,* and *ASTN1*, and with processes such as: *Potassium ion transmembrane transport, Regulation of postsynaptic membrane potential, Hormone metabolic process, Axonogenesis,* and *Synaptic transmission*. This pattern indicates the emergence of an adaptive neuroendocrine or neuroepithelial phenotype, characterized by increased functional plasticity in response to chemotherapeutic pressure^62^.

At the taxonomic dimension, the analysis of bacterial species also corroborates this functional pattern. The following were identified as central: *Bacteroides uniformis,*

*Faecalibacterium prausnitzii, Butyrivibrio crossotus, Prevotella copri, Simiaoa sunii, Vibrio harveyi,* and *Treponema succinifaciens*. Species such as *Bacteroides uniformis* and *Faecalibacterium prausnitzii* showed positive correlations with processes like response to purine-containing compound, membrane depolarization, and postsynaptic potential regulation, suggesting a direct role in the electrophysiological and metabolic modulation of the tumor epithelium^63^.

The presence of these microorganisms, especially in an environment subjected to chemotherapy, may reflect an adaptive functional recruitment with potential impacts on cell survival, microenvironment remodeling, and tumor progression. Conversely, *Butyrivibrio crossotus* demonstrated negative correlations with processes such as *regulation of epithelial cell proliferation* and *synaptic signaling*, suggesting that certain species may exert restrictive modulatory effects, possibly curbing exacerbated pro-proliferative or inflammatory pathways^64^.

Analysis of the bacterial gene–human gene–biological process axes further demonstrates that the coordinated action of bacterial genes such as *nifJ* (linked to neurogenesis and cell adhesion) and *secY/secDF* (bacterial transport and secretion) contributes to the functional and structural reorganization of the gastric epithelium. This reorganization suggests that FLOT treatment, rather than solely targeting tumor tissue, profoundly reshapes the intratumoral bacterial ecosystem, which in turn actively influences human gene expression and biological processes relevant to tumor progression and immune evasion^65^.

In summary, the microbiome of the diffuse subtype under FLOT is not a passive agent but rather an active functional modulator, capable of influencing critical pathways of tumor biology such as: Synaptic signaling and regulation of membrane potential, Ion transport and hormonal metabolism, Epithelial plasticity and cell adhesion, and Response to purinergic and sensory stimuli^66,67,68^.

Among the multiple findings of this study, the recurrent and robust correlation between the human gene *HCN1 (Hyperpolarization-activated cyclic nucleotide-gated channel 1)* and various bacterial elements, both genes and species, in the FLOT-treated diffuse subtype of GA stands out. This functional convergence positions *HCN1* as a hub for electrophysiological and signaling integration within the chemotherapy-remodeled tumor architecture, consistent with evidence indicating HCN channels as key modulators of tumor adaptation to chemotherapeutic stress through the regulation of membrane potential and cellular signaling^69^.

In the gene modulation axis, the bacterial gene *leuS*, encoding *leucyl-tRNA synthetase*, demonstrated a high-magnitude positive correlation with *HCN1*. This association is notable for two complementary reasons: (i) *leuS* is related to the maintenance of bacterial protein synthesis under stress conditions, suggesting that bacteria resilient to FLOT may activate human pathways associated with adaptive plasticity^70^; and (ii) the correlation with *HCN1* indicates that such activation may directly influence the control of epithelial membrane potential, a central factor in cellular excitability, synaptic signaling, and stress resistance^71,72^.

Complementing the modulation by bacterial genes, multiple bacterial species demonstrated a consistent correlation with *HCN1*, notably: *Bacteroides uniformis, Faecalibacterium prausnitzii, Prevotella copri, Simiaoa sunii,* and *Treponema succinifaciens.* These species represent a diverse functional spectrum: some are butyrate-producing and anti-inflammatory (e.g., *F. prausnitzii*), while others have been associated with remodeled and inflammatory tumor environments (e.g., *P. copri, Simiaoa spp.*)^73,74,75^. The convergence of these bacteria with *HCN1* reinforces the concept that FLOT treatment selects for a functional microbial consortium capable of directly interacting with the electrophysiological machinery of the tumor epithelium.

*HCN1* emerged as a functionally central gene, associated with the following enriched biological processes: *Potassium ion transmembrane transport, Regulation of membrane potential, Postsynaptic potential regulation, Modulation of chemical synaptic transmission,* and *Regulation of neuronal action potential*. The activation of these pathways in the context of gastric cancer suggests an emerging role of tumor bioelectricity as a functional axis for cellular survival and evasion of cytotoxic stress^72,76^. In models of glioblastoma, melanoma, and colon cancer, ion channels have been implicated in cellular proliferation, apoptosis, and paracrine communication within the microenvironment, mechanisms of high relevance in aggressive solid tumors like the gastric diffuse subtype^72,77,78^.

The simultaneous modulation of *HCN1* by multiple microbiome elements indicates that this gene may act as a sensor and integrator of microbial stimuli, translating environmental signals into bioelectric and functional alterations in the tumor epithelium. This finding proposes *HCN1* as a biomarker of induced bacterial functional remodeling, as well as a potential adjunct therapeutic target for patients treated with FLOT. Additionally, the association of *HCN1* with synaptic pathways and ion transport suggests that interventions targeting this channel, either pharmacologically or through microbial modulation, could alter tumor sensitivity to treatment, redefining the boundaries of personalized oncologic therapy.

## Conclusion

This study provides robust evidence that the FLOT chemotherapy regimen exerts a profound and differential remodeling effect on the gastric tumor microenvironment, integratively impacting the human transcriptome, the tumor microbiome composition, and inter-kingdom interaction networks. More than a cytotoxic agent, FLOT acts as an ecosystem modulator, capable of reconfiguring functional axes fundamental to tumor progression and therapeutic response.

In the intestinal subtype, the treatment effects reflected a coordinated activation of epithelial, inflammatory, and sensory perception pathways, involving human genes such as *CXCL8, IL1B,* and *FOSL1,* in correlation with bacterial genes like *atpD, grpE,* and *ureB,* and species such as *Lactobacillus johnsonii* and *Prevotella intermedia*. This network of interactions points to a molecularly inflamed and morphogenetically active microenvironment, possibly more responsive to therapy yet also more susceptible to plastic adaptation.

In the diffuse subtype, on the other hand, the findings revealed a more complex and adaptive pattern, with the emergence of a neuroepithelial phenotype under the direct influence of the microbiome. Bacterial genes such as *ilvD, pepN, rpoD,* and *secY,* associated with species like *Enterobacter cloacae, Klebsiella variicola,* and *Streptococcus salivarius*, showed strong correlations with human genes such as *HCN1*, *ENKUR*, *THBS1*, and *HAMP*. Among these, *HCN1* stands out as a true functional hub, integrating microbial signals with tumor electrophysiological and synaptic circuits, including processes of membrane potential modulation, cell adhesion, neural plasticity, and oxidative stress resistance. These findings suggest that the microbiome of the treated diffuse subtype not only resists FLOT but potentially co-opts it as a trigger for adaptive functional reprogramming.

In this scenario, it becomes essential to validate these findings in independent cohorts, conduct functional experimentation of the implicated pathways, and explore combined therapeutic approaches that align conventional chemotherapy with engineering of the tumor ecosystem. The integration of these elements may usher in a new generation of synergistic treatments aimed not only at cell elimination but at the intelligent reconfiguration of the tumor environment.

## Acknowledgment

The authors would like to thank the Oncology Research Center, the Human and Medical Genetics Laboratory, and the Anatomical Pathology Laboratory at João de Barros Barreto University Hospital (HUJBB – UFPA) for their invaluable technical and laboratory support. Our gratitude also goes to the High-Performance Computing Center (CCAD) at the Federal University of Pará for access to the Apollo 2000 cluster, which was crucial for our analyses. This work was supported by Fundação Amazonia de Amparo a Estudos e Pesquisas – FAPESPA (004/21), Conselho Nacional de Desenvolvimento Científico e Tecnológico – CNPq (313303/2021-5) and Ministério Público do Trabalho (11/12/2020 – Ids 372cfc4 and b7c1637).

## Data availability statement

The original contributions presented in the study are included in the article. Further inquiries can be directed to the corresponding author.

## References

1. Ilic M, Ilic I. Epidemiology of stomach cancer. World J Gastroenterol. 2022 Mar 28;28(12):1187–1203. doi: 10.3748/wjg.v28.i12.1187

2. Lin X, Yang P, Wang M, Huang X, Wang B, Chen C, Xu A, Cai J, Khan M, Liu S and Lin J (2024) Dissecting gastric cancer heterogeneity and exploring therapeutic strategies using bulk and single-cell transcriptomic analysis and experimental validation of tumor microenvironment and metabolic interplay. Front. Pharmacol. 15:1355269. doi: 10.3389/fphar.2024.1355269

3. Ma J, Shen H, Kapesa L, Zeng S. Lauren classification and individualized chemotherapy in gastric cancer. Oncol Lett. 2016 May;11(5):2959–2964. doi: 10.3892/ol.2016.4337. Epub 2016 Mar 16. PMID: 27123046; PMCID: PMC4840723.

4. Cisło M., Filip A. Anna, Arnold Offerhaus G. Johan, Ciseł B., Rawicz-Pruszyński K., Skierucha M., Polkowski W. Piotr Distinct molecular subtypes of gastric cancer: from Laurén to molecular pathology. Oncotarget. 2018; 9: 19427–19442. doi: 10.18632/oncotarget.24827.

5. Adenis A, Samalin E, Mazard T, Portales F, Mourregot A, Ychou M. Le protocole FLOT est-il le nouveau standard de chimiothérapie péri-opératoire des cancers de l’estomac[Does the FLOT regimen a new standard of perioperative chemotherapy for localized gastric cancer?]. Bull Cancer. 2020 Jan;107(1):54–60. French. doi: 10.1016/j.bulcan.2019.12.005. Epub 2020 Jan 21. PMID: 31980145.

6. Yang L, Wang Q, He L, Sun X. The critical role of tumor microbiome in cancer immunotherapy. Cancer Biol Ther. 2024 Dec 31;25(1):2301801. doi: 10.1080/15384047.2024.2301801. Epub 2024 Jan 19. PMID: 38241173; PMCID: PMC10802201.

7. Boesch M, Horvath L, Baty F, Pircher A, Wolf D, Spahn S, Straussman R, Tilg H, Brutsche MH. Compartmentalization of the host microbiome: how tumor microbiota shapes checkpoint immunotherapy outcome and offers therapeutic prospects. J Immunother Cancer. 2022 Nov;10(11):e005401. doi: 10.1136/jitc-2022-005401.

8. Geller LT, Barzily-Rokni M, Danino T, Jonas OH, Shental N, Nejman D, Gavert N, Zwang Y, Cooper ZA, Shee K, Thaiss CA, Reuben A, Livny J, Avraham R, Frederick DT, Ligorio M, Chatman K, Johnston SE, Mosher CM, Brandis A, Fuks G, Gurbatri C, Gopalakrishnan V, Kim M, Hurd MW, Katz M, Fleming J, Maitra A, Smith DA, Skalak M, Bu J, Michaud M, Trauger SA, Barshack I, Golan T, Sandbank J, Flaherty KT, Mandinova A, Garrett WS, Thayer SP, Ferrone CR, Huttenhower C, Bhatia SN, Gevers D, Wargo JA, Golub TR, Straussman R. Potential role of intratumor bacteria in mediating tumor resistance to the chemotherapeutic drug gemcitabine. Science. 2017 Sep 15;357(6356):1156-1160. doi: 10.1126/science.aah5043.

9. Xavier JB, Young VB, Skufca J, Ginty F, Testerman T, Pearson AT, Macklin P, Mitchell A, Shmulevich I, Xie L, Caporaso JG, Crandall KA, Simone NL, Godoy-Vitorino F, Griffin TJ, Whiteson KL, Gustafson HH, Slade DJ, Schmidt TM, Walther-Antonio MRS, Korem T, Webb-Robertson BM, Styczynski MP, Johnson WE, Jobin C, Ridlon JM, Koh AY, Yu M, Kelly L, Wargo JA. The Cancer Microbiome: Distinguishing Direct and Indirect Effects Requires a Systemic View. Trends Cancer. 2020 Mar;6(3):192–204. doi: 10.1016/j.trecan.2020.01.004. Epub 2020 Feb 7. PMID: 32101723; PMCID: PMC7098063.

10. Žukauskaitė K, Baušys B, Horvath A, Sabaliauskaitė R, Šeštokaitė A, Mlynska A, Jarmalaitė S, Stadlbauer V, Baušys R, Baušys A. Gut Microbiome Changes After Neoadjuvant Chemotherapy and Surgery in Patients with Gastric Cancer. Cancers (Basel). 2024 Dec 5;16(23):4074. doi: 10.3390/cancers16234074. PMID: 39682264; PMCID: PMC11640656.

11. Li N, Bai C, Zhao L, Sun Z, Ge Y, Li X. The Relationship Between Gut Microbiome Features and Chemotherapy Response in Gastrointestinal Cancer. Front Oncol. 2021 Dec 23;11:781697. doi: 10.3389/fonc.2021.781697. PMID: 35004303; PMCID: PMC8733568.

12. Wood, D.E., Lu, J. & Langmead, B. Improved metagenomic analysis with Kraken 2. Genome Biol 20, 257 (2019). doi: 10.1186/s13059-019-1891-0.

13. Patro, R., Duggal, G., Love, M. et al. Salmon provides fast and bias-aware quantification of transcript expression. Nat Methods 14, 417–419 (2017). doi: 10.1038/nmeth.4197.

14. Sayers, E. W. et al. Database resources of the National Center for Biotechnology Information. Nucleic Acids Res. 50, D20–D26 (2022). doi: 10.1093/nar/gkab1112.

15. Frankish, A. et al. GENCODE reference annotation for the human and mouse genomes. Nucleic Acids Res. 47, D766–D773 (2019). 10.1093/nar/gky955.

16. Love, M.I., Huber, W. & Anders, S. Moderated estimation of fold change and dispersion for RNA-seq data with DESeq2. Genome Biol 15, 550 (2014). 10.1186/s13059-014-0550-8

17. Jombart, T., Devillard, S. & Balloux, F. Discriminant analysis of principal components: a new method for the analysis of genetically structured populations. BMC Genet 11, 94 (2010). 10.1186/1471-2156-11-94

18. Yu, G., Wang, L.-G., Han, Y. & He, Q.-Y. clusterProfiler: an R package for comparing biological themes among gene clusters. OMICS 16, 284–287 (2012). 10.1089/omi.2011.0118

19. Ashburner, M. et al. Gene ontology: tool for the unification of biology. Nat. Genet. 25, 25–29 (2000). 10.1038/75556

20. Carlson, M. org.Hs.eg.db: Genome Wide Annotation for Human. R package version X.Y.Z. (2023). https://bioconductor.org/packages/org.Hs.eg.db

21. Wei, T. & Simko, V. corrplot: Visualization of a Correlation Matrix. R package version 0.92 (2021). https://github.com/taiyun/corrplot

22. Brunson, J. ggalluvial: Alluvial Diagrams in ‘ggplot2’. R package version 0.12.3 (2023). https://cran.r-project.org/package=ggalluvial

23. Liang J, Liu J, Deng Z, Liu Z, Liang L. DLX6 promotes cell proliferation and survival in oral squamous cell carcinoma. Oral Dis. 2022 Jan;28(1):87–96. doi: 10.1111/odi.13728. Epub 2020 Dec 14. PMID: 33215805.

24. Han, Z.-J., Li, Y.-B., Yang, L.-X., Cheng, H.-J., Liu, X., & Chen, H. (2022). Roles of the CXCL8-CXCR1/2 Axis in the Tumor Microenvironment and Immunotherapy. Molecules, 27(1), 137. 10.3390/molecules27010137

25. Shetty, A., Tripathi, S. K., Junttila, S., Buchacher, T., Biradar, R., Bhosale, S. D., Envall, T., Laiho, A., Moulder, R., Rasool, O., Galande, S., Elo, L. L., & Lahesmaa, R. (2022). A systematic comparison of FOSL1, FOSL2 and BATF-mediated transcriptional regulation during early human Th17 differentiation. Nucleic acids research, 50(9), 4938–4958. 10.1093/nar/gkac256

26. Wang, B., Li, X., & Gao, X. (2025). Meteorin-β: A Novel Biomarker and Therapeutic Target on Its Way to the Regulation of Human Diseases. International journal of molecular sciences, 26(10), 4485. 10.3390/ijms26104485

27. Simonson, L., Vold, S., Mowers, C., Massey, R. J., Ong, I. M., Longley, B. J., & Chang, H. (2020). Keratin 13 deficiency causes white sponge nevus in mice. Developmental biology, 468(1-2), 146–153. 10.1016/j.ydbio.2020.07.016

28. Xue, X., Guo, Y., Zhao, Q., Li, Y., Rao, M., Qi, W., & Shi, H. (2023). Weighted Gene Co-Expression Network Analysis of Oxymatrine in Psoriasis Treatment. Journal of inflammation research, 16, 845–859. 10.2147/JIR.S402535

29. Li, M., Li, H., Yuan, T., Liu, Z., Li, Y., Tan, Y., & Long, Y. (2024). MUC21: a new target for tumor treatment. Frontiers in oncology, 14, 1410761. 10.3389/fonc.2024.1410761

30. Lee, Dh., Ahn, H., Sim, HI. et al. A CRISPR activation screen identifies MUC-21 as critical for resistance to NK and T cell-mediated cytotoxicity. J Exp Clin Cancer Res 42, 272 (2023). 10.1186/s13046-023-02840-9

31. Esberg, A., Haworth, S., Brunius, C. et al. Carbonic Anhydrase 6 Gene Variation influences Oral Microbiota Composition and Caries Risk in Swedish adolescents. Sci Rep 9, 452 (2019). 10.1038/s41598-018-36832-z.

32. Mboge, M. Y., Mahon, B. P., McKenna, R., & Frost, S. C. (2018). Carbonic Anhydrases: Role in pH Control and Cancer. Metabolites, 8(1), 19. 10.3390/metabo8010019.

33. Browning K. N. (2015). Role of central vagal 5-HT3 receptors in gastrointestinal physiology and pathophysiology. Frontiers in neuroscience, 9, 413. 10.3389/fnins.2015.00413.

34. Lu X, Xu H, Xu J, Lu S, You S, Huang X, Zhang N and Zhang L (2022) The regulatory roles of the E3 ubiquitin ligase NEDD4 family in DNA damage response. Front. Physiol. 13:968927. doi: 10.3389/fphys.2022.968927.

35. Rotin D and Staub O (2012) Nedd4-2 and the regulation of epithelial sodium transport. Front. Physio. 3:212. doi: 10.3389/fphys.2012.00212.

36. Song, C., Orlandi, C., Sutton, L. P., & Martemyanov, K. A. (2019). The signaling proteins GPR158 and RGS7 modulate excitability of L2/3 pyramidal neurons and control A-type potassium channel in the prelimbic cortex. The Journal of biological chemistry, 294(35), 13145–13157. 10.1074/jbc.RA119.007533.

37. Iden, J. A., Raphael-Mizrahi, B., Awida, Z., Naim, A., Zyc, D., Liron, T., Kasher, M., Livshits, G., Vered, M., & Gabet, Y. (2023). The Anti-Tumorigenic Role of Cannabinoid Receptor 2 in Colon Cancer: A Study in Mice and Humans. International Journal of Molecular Sciences, 24(4), 4060. 10.3390/ijms24044060

38. Elbaz M., Ahirwar D., Ravi J., Nasser M. W., Ganju R. K. Novel role of cannabinoid receptor 2 in inhibiting EGF/EGFR and IGF-I/IGF-IR pathways in breast cancer. Oncotarget. 2017; 8: 29668–29678. Retrieved from https://www.oncotarget.com/article/9408/text/

39. Zu, M., Guo, WW., Cong, T. et al. SCN11A gene deletion causes sensorineural hearing loss by impairing the ribbon synapses and auditory nerves. BMC Neurosci 22, 18 (2021). 10.1186/s12868-021-00613-8.

40. Duvivier, L., Gerard, L., Diaz, A., & Gillet, J.-P. (2024). Linking ABC transporters to the hallmarks of cancer. Trends in Cancer, 10(2), 124–134. 10.1016/j.trecan.2023.09.013.

41. Wang, J., Yunyun, Z., Wang, L., Chen, X., & Zhu, Z. (2017). ABCG2 confers promotion in gastric cancer through modulating downstream CRKL in vitro combining with biostatistics mining. Oncotarget, 8(3), 5256–5267. 10.18632/oncotarget.14128.

42. Krawiec P, Pac-Kozuchowska E. Rola hepcydyny w metabolizmie żelaza w przebiegu nieswoistych zapaleń jelit [The role of hepcidin in iron metabolism in inflammatory bowel diseases]. Postepy Hig Med Dosw (Online). 2014;68:936–43. Polish. doi: 10.5604/17322693.1111365. PMID: 24988613.

43. Ma, Q., Lu, Y., Lin, J., & Gu, Y. (2019). ENKUR acts as a tumor suppressor in lung adenocarcinoma cells through PI3K/Akt and MAPK/ERK signaling pathways. Journal of Cancer, 10(17), 3975–3984. 10.7150/jca.30021.

44. Ma, Q., Lu, Y., & Gu, Y. (2019). ENKUR Is Involved in the Regulation of Cellular Biology in Colorectal Cancer Cells via PI3K/Akt Signaling Pathway. Technology in cancer research & treatment, 18, 1533033819841433. 10.1177/1533033819841433

45. Rah B, Farhat NM, Hamad M, Muhammad JS. JAK/STAT signaling and cellular iron metabolism in hepatocellular carcinoma: therapeutic implications. Clin Exp Med. 2023 Nov;23(7):3147–3157. doi: 10.1007/s10238-023-01047-8. Epub 2023 Mar 28. PMID: 36976378.

46. Kowdley, K. V., Gochanour, E. M., Sundaram, V., Shah, R. A., & Handa, P. (2021). Hepcidin Signaling in Health and Disease: Ironing Out the Details. Hepatology communications, 5(5), 723–735. 10.1002/hep4.1717.

47. Xu, J., Jin, Y., Wang, M., Tao, Y., Wang, Y., & Gong, F. (2025). Amphiregulin Promotes Proliferation and Migration of the Damaged Endothelial Cells in Kawasaki Disease Cell Models. Immunity, inflammation and disease, 13(7), e70223. 10.1002/iid3.70223.

48. Hua, F., & Tian, Y. (2017). CCL4 promotes the cell proliferation, invasion and migration of endometrial carcinoma by targeting the VEGF-A signal pathway. International journal of clinical and experimental pathology, 10(11), 11288–11299.

49. Guan, Y.-T., Zhang, C., Zhang, H.-Y., Wei, W.-L., Yue, W., Zhao, W., & Zhang, D.-H. (2023). Primary cilia: Structure, dynamics, and roles in cancer cells and tumor microenvironment. Journal of Cellular Physiology, 238, 1788–1807. 10.1002/jcp.31092

50. Fabbri, L., Bost, F., & Mazure, N. M. (2019). Primary Cilium in Cancer Hallmarks. International Journal of Molecular Sciences, 20(6), 1336. 10.3390/ijms20061336.

51. Rébé, C., & Ghiringhelli, F. (2020). Interleukin-1β and Cancer. Cancers, 12(7), 1791. 10.3390/cancers12071791.

52. Plati, J., Bucur, O., & Khosravi-Far, R. (2011). Apoptotic cell signaling in cancer progression and therapy. Integrative biology: quantitative biosciences from nano to macro, 3(4), 279–296. 10.1039/c0ib00144a.

53. Tong, Y., Gao, H., Qi, Q., Liu, X., Li, J., Gao, J., Li, P., Wang, Y., Du, L., & Wang, C. (2021). High fat diet, gut microbiome and gastrointestinal cancer. Theranostics, 11(12), 5889–5910. 10.7150/thno.56157.

54. Chen Z, Guan D, Wang Z, et al. Microbiota in cancer: molecular mechanisms and therapeutic interventions. MedComm. 2023; 4:e417. 10.1002/mco2.417.

55. Heidari, M., Maleki Vareki, S., Yaghobi, R., & Karimi, M. H. (2024). Microbiota activation and regulation of adaptive immunity. Frontiers in immunology, 15, 1429436. 10.3389/fimmu.2024.1429436.

56. von Frieling, J., Fink, C., Hamm, J., Klischies, K., Forster, M., Bosch, T. C. G., Roeder, T., Rosenstiel, P., & Sommer, F. (2018). Grow With the Challenge – Microbial Effects on Epithelial Proliferation, Carcinogenesis, and Cancer Therapy. Frontiers in microbiology, 9, 2020. 10.3389/fmicb.2018.02020.

57. Bose, S., Banerjee, S., Mondal, A., Chakraborty, U., Pumarol, J., Croley, C. R., & Bishayee, A. (2020). Targeting the JAK/STAT Signaling Pathway Using Phytocompounds for Cancer Prevention and Therapy. Cells, 9(6), 1451. 10.3390/cells9061451.

58. Li, S., Zhu, S., & Yu, J. (2024). The role of gut microbiota and metabolites in cancer chemotherapy. Journal of advanced research, 64, 223–235. 10.1016/j.jare.2023.11.027.

59. Ma W, Mao Q, Xia W, Dong G, Yu C and Jiang F (2019) Gut Microbiota Shapes the Efficiency of Cancer Therapy. Front. Microbiol. 10:1050. doi: 10.3389/fmicb.2019.01050.

60. Loh, J.S., Mak, W.Q., Tan, L.K.S. et al. Microbiota–gut–brain axis and its therapeutic applications in neurodegenerative diseases. Sig Transduct Target Ther 9, 37 (2024). 10.1038/s41392-024-01743-1.

61. Murciano-Brea, J., Garcia-Montes, M., Geuna, S., & Herrera-Rincon, C. (2021). Gut Microbiota and Neuroplasticity. Cells, 10(8), 2084. 10.3390/cells10082084.

62. Ciernikova, S., Mego, M., & Chovanec, M. (2021). Exploring the Potential Role of the Gut Microbiome in Chemotherapy-Induced Neurocognitive Disorders and Cardiovascular Toxicity. Cancers, 13(4), 782. 10.3390/cancers13040782.

63. Mahapatra, C., Kishore, A., Gawad, J., Al-Emam, A., Kouzeiha, R. A., & Rusho, M. A. (2025). Review of electrophysiological models to study membrane potential changes in breast cancer cell transformation and tumor progression. Frontiers in physiology, 16, 1536165. 10.3389/fphys.2025.1536165.

64. Ionica, E., Gaina, G., Tica, M., Chifiriuc, M. C., & Gradisteanu-Pircalabioru, G. (2022). Contribution of Epithelial and Gut Microbiome Inflammatory Biomarkers to the Improvement of Colorectal Cancer Patients’ Stratification. Frontiers in oncology, 11, 811486. 10.3389/fonc.2021.811486.

65. Zhao, LY., Mei, JX., Yu, G. et al. Role of the gut microbiota in anticancer therapy: from molecular mechanisms to clinical applications. Sig Transduct Target Ther 8, 201 (2023). 10.1038/s41392-023-01406-7.

66. Glinert, A., Turjeman, S., Elliott, E., & Koren, O. (2022). Microbes, metabolites and (synaptic) malleability, oh my! The effect of the microbiome on synaptic plasticity. Biological reviews of the Cambridge Philosophical Society, 97(2), 582–599. 10.1111/brv.12812.

67. Silva YP, Bernardi A and Frozza RL (2020) The Role of Short-Chain Fatty Acids From Gut Microbiota in Gut-Brain Communication. Front. Endocrinol. 11:25. doi: 10.3389/fendo.2020.00025.

68. Roy, R., & Singh, S. K. (2024). The Microbiome Modulates the Immune System to Influence Cancer Therapy. Cancers, 16(4), 779. 10.3390/cancers16040779.

69. Zhao J, Li M, Xu J and Cheng W (2022) The modulation of ion channels in cancer chemo-resistance. Front. Oncol. 12:945896. doi: 10.3389/fonc.2022.945896.

70. Karkhanis, V. A., Mascarenhas, A. P., & Martinis, S. A. (2007). Amino acid toxicities of Escherichia coli that are prevented by leucyl-tRNA synthetase amino acid editing. Journal of bacteriology, 189(23), 8765–8768. 10.1128/JB.01215-07.

71. Huang, Z., Walker, M. C., & Shah, M. M. (2009). Loss of dendritic HCN1 subunits enhances cortical excitability and epileptogenesis. The Journal of neuroscience: the official journal of the Society for Neuroscience, 29(35), 10979–10988. 10.1523/JNEUROSCI.1531-09.2009

72. Mok, K. C., Tsoi, H., Man, E. P., Leung, M. H., Chau, K. M., Wong, L. S., Chan, W. L., Chan, S. Y., Luk, M. Y., Chan, J. Y. W., Leung, J. K. M., Chan, Y. H. Y., Batalha, S., Lau, V., Siu, D. C. W., Lee, T. K. W., Gong, C., & Khoo, U. S. (2021). Repurposing hyperpolarization-activated cyclic nucleotide-gated channels as a novel therapy for breast cancer. Clinical and translational medicine, 11(11), e578. 10.1002/ctm2.578

73 Dikeocha, I. J., Al-Kabsi, A. M., Chiu, H. T., & Alshawsh, M. A.. (2022). Faecalibacterium prausnitzii Ameliorates Colorectal Tumorigenesis and Suppresses Proliferation of HCT116 Colorectal Cancer Cells. Biomedicines, 10(5), 1128. 10.3390/biomedicines10051128

74 Obuya S, Elkholy A, Avuthu N, Behring M, Bajpai P, Agarwal S, Kim HG, El-Nikhely N, Akinyi P, Orwa J, Afaq F, Abdalla M, Michael A, Farouk M, Bateman LB, Fouad M, Saleh M, Guda C, Manne U, Arafat W. A signature of Prevotella copri and Faecalibacterium prausnitzii depletion, and a link with bacterial glutamate degradation in the Kenyan colorectal cancer patients. J Gastrointest Oncol 2022;13(5):2282–2292. doi: 10.21037/jgo-22-116

75 Newman, T.M., Shively, C.A., Register, T.C., et al. Diet, obesity, and the gut microbiome as determinants modulating metabolic outcomes in a non-human primate model. Microbiome 9, 100 (2021). 10.1186/s40168-021-01069-y

76 Sheth M and Esfandiari L (2022) Bioelectric Dysregulation in Cancer Initiation, Promotion, and Progression. Front. Oncol. 12:846917. doi: 10.3389/fonc.2022.846917

77. Gentile, R., Feudi, D., Sallicandro, L., & Biagini, A. (2025). Can the Tumor Microenvironment Alter Ion Channels? Unraveling Their Role in Cancer. Cancers, 17(7), 1244. 10.3390/cancers17071244

78 Liang, X., Zhang, J., & Luo, Y. (2024). Investigating the role of hyperexpressed HCN1 in inducing myocardial infarction through activation of the NF-κB signaling pathway. Open life sciences, 19(1), 20220967. 10.1515/biol-2022-0967

